# KBPRNA: A novel method integrating bulk RNA-seq data and LINCS-L1000 gene signatures to predict kinase activity based on machine learning

**DOI:** 10.1101/2022.11.16.516707

**Authors:** Yuntian Zhang, Lantian Yao, Yixian Huang, Wenyang Zhang, Yuxuan Pang, Tzongyi Lee

**Affiliations:** Warshel institute for computational Biology, The Chinese university of HongKong, Shenzhen; School of Life and Health science, School of Medicine, The Chinese university of HongKong, Shenzhen; Kobilka Institute of Innovative Drug Discovery, The Chinese university of HongKong, Shenzhen; School of science and Engineering, The Chinese university of HongKong, Shenzhen

**Keywords:** Kinase activity, bulk RNA-sequence technology, LINCS-L1000, XGBoost algorithm

## Abstract

**Background:** Kinases are a type of enzymes which can transfer phosphate groups from high-energy and phosphate-donating molecules to specific substrates. Kinase activities could be utilized to be represented as specific biomarkers of specific cancer types. Nowadays novel algorithms have already been developed to compute kinase activities from phosphorylated proteomics data. However, phosphorylated proteomics sequencing could be costly expensive and need valuable samples. Moreover,not methods which could achieve kinase activities from bulk RNA-sequence data have been developed. Here we propose KBPRNA, a general computational framework for extracting specific kinase activities from bulk RNA-sequencing data in cancer samples. KBPRNA also achieves better performance in predicting kinase activities from bulk RNA-sequence data under cancer conditions benchmarking against other models.

**Results:** In this study, we used LINCS-L1000 dataset which was used to be reported as efficient gene signatures in defining bulk RNA-seq data as input dataset of KBPRNA. Also, we utilized eXtreme Gradient Boosting (XGboost) as the main algorithm to extract valuable information to predict kinase activities. This model outperforms other methods such as linear regression and random forest in predicting kinase activities from bulk RNA-seq data. KBPRNA integrated tissue samples coming from breast invasive carcinoma, hepatocellular carcinoma, lung squamous cell carcinoma, Glioblastoma multiforme and Uterine Corpus Endometrial Carcinoma. It was found that KBPRNA achieved good performance with an average R score above threshold of 0.5 in kinase activity prediction.

**Conclusions:** Model training and testing process showed that KBPRNA outperformed other machine learning methods in predicting kinase activities coming from various cancer types’ tissue samples. This model could be utilized to approximate basic kinase activities and link it with specific biological functions, which in further promoted the progress of cancer identification and prognosis.

## Background

Kinases are a type of enzymes that catalyzes the transfer of phosphate groups from high-energy, phosphate-donating molecules to specific substrates [1]. Kinases play a great role in the pathogenesis of cancer and several kinases have been investigated as drug targets [2]. Multitargeted receptor tyrosine kinase (RTK) inhibitors have been approved as the specific medicine for the cancer [3]. Until 2020, 52 small molecule protein kinase inhibitors have been approved by the US FDA [4]. Thus, understanding kinase activity profiles in cancer tissues is fundamental for the treatment of cancer. However, most kinases have not been well studied and the discovery of kinase inhibitors still has a long way to go [5].

Traditionally, the measurement of protein kinase activity relied on the use of radioactivity and incorporation of 32P into proteins or peptide substrates [6, 7]. A disadvantage of these methods is that they are not suitable for measuring the kinase activity of a large number of kinases at the same time. More recently, these methods have been optimized for high-throughput applications with the general goal of testing for inhibitor specificity [8]. Because of the negative implications of the use of radioactivity, non-radioactive methods for measuring kinase activity have become increasingly popular. Many of these methods are based on the measurement of fluorescent peptide substrates and take advantage of altered fluorescent properties upon phosphorylation [9]. Furthermore, some of these fluorescent- and luminescent-based methods are amenable to studies in cell lysates and live cells, providing a more natural environment for studying kinase activity. However, the cost of these methods in measuring kinase activities is still high and there are large amounts of challenges in measuring a wide range of kinase activities.

With the maturity of high-throughput sequencing technology, proteomics sequencing data could be exploited to compute corresponding kinases’ activities [10]. Multiple computational tools which could be applied to predict kinase inhibitor resistance and selectivity have already been developed [11, 12]. Emilio Fenoy et al developed a generic deep convolutional neural network framework called NetPhosPan to kinase phosphorylation prediction [13]. Kathryn E Kirchoff et al proposed a deep learning model called EMBER for multi-label prediction of kinase-substrate phosphorylation events [14]. Several studies have already reported that substrate genes’ RNA-seq expression and kinase activities under cancer conditions could be related [15]. However, no study has demonstrated the specific relationships between kinase activities and RNA-seq expression level in cancer tissues. No study has covered whether specific kinase activities could be used as biomarkers for distinguishing different cancer types.

In this research, we proposed an interpretable machine learning model KBPRNA based on bulk RNA-seq data from various cancer types to predict kinase activities. LINCS-L1000 gene signatures were integrated for feature selection. A modified XGBoost model was utilized as the basic algorithm of this method. By five-fold cross-validation approach, we evaluated KBPRNA using different feature sets and across various machine learning algorithms and parameters. KBPRNA which combines both LINCS-L1000 gene signatures and XGBoost model shows good performance based on bulk RNA-seq data collected from independent cancer samples [16–20]. After getting kinases with high predictability based on bulk RNA-seq data, we utilized machine learning model to predict specific cancer types based on specific kinase activities’ profiles. It was found that specific kinase activities could be used as input features to classify different cancer types. KBPRNA was also applied to scRNA-seq dataset of breast invasive carcinoma tissues to achieve each individual cell’s kinase activity profiles based on each individual cell’s RNA-seq profiles. Then kinase activity profiles were constructed according to cell groups identified through a scRNA-seq data analysis tool scanpy [21]. Kinases which can differentiate multiple cell groups were identified in this process. In sum, this research shows the ability of specific kinase activities in cancer type identification and scRNA-seq cell group clustering.

## Methods

All the analysis code that we implemented was executed in Python version 3.8.1. We carried all the statistical tests using the R computing environment (version 4.0.0). Flowchart was drawn using biorender [22]. The rest pictures were drawn using Matplotlib package in Python version 3.8.1.

### Data Collection and preprocessing

Bulk RNA-seq data collection: 5 types’ cancer samples’ bulk RNA-seq dataset and corresponding phosphorylated proteomics dataset which are Breast cancer, Glioblastroma multiforme, Hepatocellular Carcinoma, lung squamous cell carcinoma, endometrial carcinoma are collected from supplementary files of recent journals [23]. Then a computing method named KSEA was utilized to compute kinase activities from five cancer samples’ phosphorylated proteomics dataset [24]. Computed kinase activity profiles have been deposited in github: https://github.com/tibettiger/kinase_prediction/tree/main/data/kinase_activity. scRNA-seq data collection: scRNA-seq dataset of breast invasive carcinoma: GSM5457199 was selected as one application sample of KBPRNA to construct single cell kinase activity profiles [25].

### Model establishment

LINC-L1000 dataset is a set of gene signatures which can be utilized to represent the whole transcriptomes of humans. LINC-L1000 dataset has already been proved to be efficient in predicting drug-induced cell viability and drug-drug interaction (DDI) [26]. LINCS L1000 is generated using Luminex L1000 technology where the expression levels of 978 landmark genes are measured by fluorescence intensity. This research shows that LINC-L1000 dataset outperformed other gene signatures in predicting kinase activity under cancer circumstances. KBPRNA model utilized to predict kinase activity under specific cancer conditions is composed of two parts. Firstly, whole transcriptomics data was transformed into dataset comprised of only LINCS-L1000 gene signatures and genes which don’t belong to LINCS-L1000 gene signatures were excluded. Secondly, XGBoost which proved efficient in predicting both continuous or discrete data was applied to predict kinase activity based on LINCS-L1000-transformed gene expression dataset [27]. A prediction score on the leaf node of each decision tree based on the differential expression of genes in each sample was obtained and multiple weak estimators are constructed one by one through multiple iterations. The kinase activity prediction result is defined as the sum of the prediction scores of all the trees as follows:

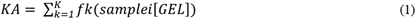

where *KA* represents kinase activity levels, *fk*(*samplei*[*GEL*]) represents the prediction score on the *k*-th decision tree for the *i*-th sample on the LINCS-L1000 transformed gene expression levels. *K* is the number of decision trees. Then during the *t*-th iteration of the sample, the model’s predicted value *KA* can be described as follows:

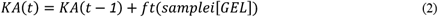

In order to reduce the prediction bias and improve the prediction accuracy, 5-fold cross-validation using *GridSearchCv* method from python package scikit-learn version 0.19.1 was utilized to find parameters which show best performance to predict the corresponding kinase activity based on bulk RNA-seq data coming from cancer samples [28].

The adjustable parameters of XGBoost model include: 1) max_depth = [4,5,6,7] 2) learning_rate =[0.03,0.06,0.09,0.12,0.15,0.18,0.21,0.24,0.26,0.3] 3) n_estimators = [100,200]

In order to measure the efficiency of this model which combines both XGBoost model and LINCS-L1000 gene signatures, this method was benchmarked against three combinations: LINCS-L1000 gene signatures + Linear regression model; All gene signatures + XGBoost model; All gene signatures + Linear regression model.

The above method was applied to five type cancer samples (Breast cancer, Glioblastroma multiforme, Hepatocellular Carcinoma, lung squamous cell carcinoma, endometrial carcinoma) and predict corresponding kinase activity for each cancer type. R square was utilized to measure the performance of the model in predicting kinase activity and R square formula is shown as below:

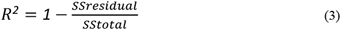

where *SSresidual* represents residual sum of squares, *SStotal* represents total sum of squares.

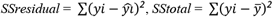

where *yi* represents actual number of targets, 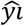 represents predicted number of targets and *ӯ* represents average number of all targets.

### Cancer type classification based on specific kinases

After getting the top 50 kinases which show its performance in representing RNA-seq, a classifier using RandomForest, XGBoost or other ensemble learning methods to predict each samples’ cancer type was constructed. AUC curve was utilized to demonstrate the performance of the classifier. We made a list which ranked all kinds of kinases’ predictability. Top10 kinases were chosen from the list and low10 kinases from the list were also selected as the input features of our classifier in order to demonstrate the superiority of top10 kinases with highest predictability in classifying specific cancer type.

### Single cell kinase profiles construction based on KBPRNA

In order to demonstrate the ability of top 50 kinases as the biomarkers for distinguishing different cell groups from one sample, kinase activity profiles of each cell from breast cancer sample were constructed using our pretrained model which involve 50 kinases [18]. Then scanpy was utilized to cluster several cell groups from scRNA-seq datasets of breast cancer tissues. Our results show similar clustering performance compared to that when using real scRNA-seq datasets [29].

## Results

### Pearson correlations between kinase activities and substrates’ gene expression

To investigate pearson correlation between kinase activities and substrates’ gene expression, we compute kinase activities using KSEA algorithm and utilize our previous kinase-substrate database to measure the pearson correlations between kinase activities and substrates’ gene expression. Under different cancer conditions, we notice that most kinase activities do not perform a high correlation with substrates’ gene expression. Under hepatocellular carcinoma, kinase TNK2 shows significantly negative correlated with CAT gene expression (PCC = −0.8). On the contrary, kinase TNK2 shows significantly positive correlated with CAT gene expression (PCC = 0.89). This situation also happened to other kinase-substrate pairs, such as ARAF-CAT pairs under hepatocellular carcinoma (PCC = −0.81) and lung squamous cell carcinoma (PCC = −0.87). At the same time, some kinase/substrate pairs show consistent associations under two different cancer conditions. For instance, ABL1 activity values shows negative correlated with CAT gene expression under both hepatocellular carcinoma and lung squamous cell carcinoma. In sum, most kinase/substrate pairs do not show obvious connections in terms of Pearson correlations. Also, kinase activity does not perform consistent impact on substrates’ gene expression such as inhibition or promotion under various cancer conditions. Above all, inconsistent Pearson correlations between kinase activities and substrates’ gene expression indicate that the relationships between kinase activities and substrates’ gene expression are complex and may vary with cancer types. Thus, we need novel machine learning method for us to estimate specific kinase activities instead of single substrates’ gene expression.

### Performance of LINCS-L1000 genes plus XGBoost versus all genes or linear regression

The LINCS-L1000 dataset is a comprehensive resource for gene expression changes observed in human cell lines perturbed with small molecules and genetic constructs [30]. It utilized 978 genes to extrapolate the expression of the whole genome. Here, we proposed a novel machine learning method which combines both LINCS-L1000 genes and XGBoost. XGBoost is one of the most advantageous prediction algorithms and could have high performance in both classification and regression tasks [31]. Here, we used XGBoost algorithm to predict kinase activities and predicted values are obtained from the leaf node of each decision tree based on the differential expression of genes in each sample. Multiple iterations construct multiple weak estimators. Kinase activities prediction result is defined as the sum of the prediction scores of all the trees as follows.

In this study, in order to improve the prediction accuracy of kinase activities and minimize the prediction bias, we used Gridsearch and cross validation to find the best parameters of XGBoost model. We set adjustable parameters of XGBoost as follows: max_depth = [4, 5, 6], learning_rate =(0.03, 0.06, 0.09, 0.12, 0.15, 0.18, 0.21, 0.24, 0.27, 0.3), n_estimators =[100, 200]. Finally, we used R square to measure the performance of our model.

We selected seven representative kinases which show wide distributions among cancer tissues respectively from five types’ cancer: breast invasive carcinoma (CDK2, ERN1, CIT, TNK2, CDK4, MKK2, CDK5); hepatocellular carcinoma (AOK3, iRPK1, MKK2, CIT, P4K5, YES1, RAK1); lung squamous cell carcinoma (ERN1, CDK1, PLK1, NK1E, ARAF, TNK2, iRPK1); glioblastoma multiforme (M20C, MINK1, IRPK1, TNK2, EGFR, CIT); uterine corpus endometrial carcinoma (PLK1, ARAF, IP4K5, IP3K1, NK1E, ERN1, IP4K4). Among all the four methods of predicting kinase activities (L1000 + XGBoost; L1000 + Linear_regression; Allgene +XGboost; Allgene +Linear_regression), method “L1000 +XGBoost” performs best in all the five cancer types. This model achieves better performance even in dataset of glioblastoma multiforme. These results demonstrate advantageous of our model. The prediction results and corresponding XGBoost model’s parameters are shown in supplementary table S1.

After building all the models of kinase prediction using bulk RNA-seq dataset, we noticed that this model could predict over 50% kinases’ activities accurately (R square > 0.5) under conditions of hepatocellular carcinoma and lung squamous cell carcinoma. However, under conditions of glioblastoma multiforme and breast invasive carcinoma, this model does not perform as well as it does under conditions of hepatocellular carcinoma and lung squamous cell carcinoma. This phenomenon may result from cancer type heterogeneity and batch effect of each sample. It also denotes that the same kinases’ predictability may vary with cancer types.

### Cancer type identification based on kinases with high predictability

In order to demonstrate this model’s robustness, we rank the common 120 kinases coming from tissues of cancer types according to their predictability (supplementary tableS2). Ten kinases’ activities were selected iteratively from the list. The top 10 kinases represent kinases with the most predictability. At the same time, the bottom 10 kinases represent kinases with the least predictability. We utilized ten kinases from the list as the input feature of our XGBoost model to predict the sample’s corresponding cancer type. Total 866 samples were input into this cancer type prediction model. Our model achieves perfect performance in cancer type classification using top10 kinases as our input features (error rate = 4.608%, Fig4B). We benchmarked this result against other method using ten kinases with lower predictability (Fig4C). The classification accuracy of each ten kinases as input features coincidentally corresponded to our previous list of kinase predictability. Top10 kinases achieved highest classification accuracy in differentiating cancer types (Accuracy = 95.39%). Term “21-30 kinases”, “41-50 kinases”, “71-80 kinases”, “91-100 kinases” followed it with classification accuracy respectively (accuracy = 88.48%, 88.94%, 77.42%, 74.20%). Low10 kinases achieved lowest classification accuracy in differentiate different cancer types (Accuracy = 60.83%). ROC curve was also used to compare the cancer type classification performance between top10 kinases and low10 kinases (Fig4D). Top10 kinases as the input features achieved AUC area equaling to 0.75, larger than AUC area equaling 0.43 using low10 kinases as the input features. These results demonstrate superiority of our model in identifying kinases which can own high predictability and classify specific cancer types.

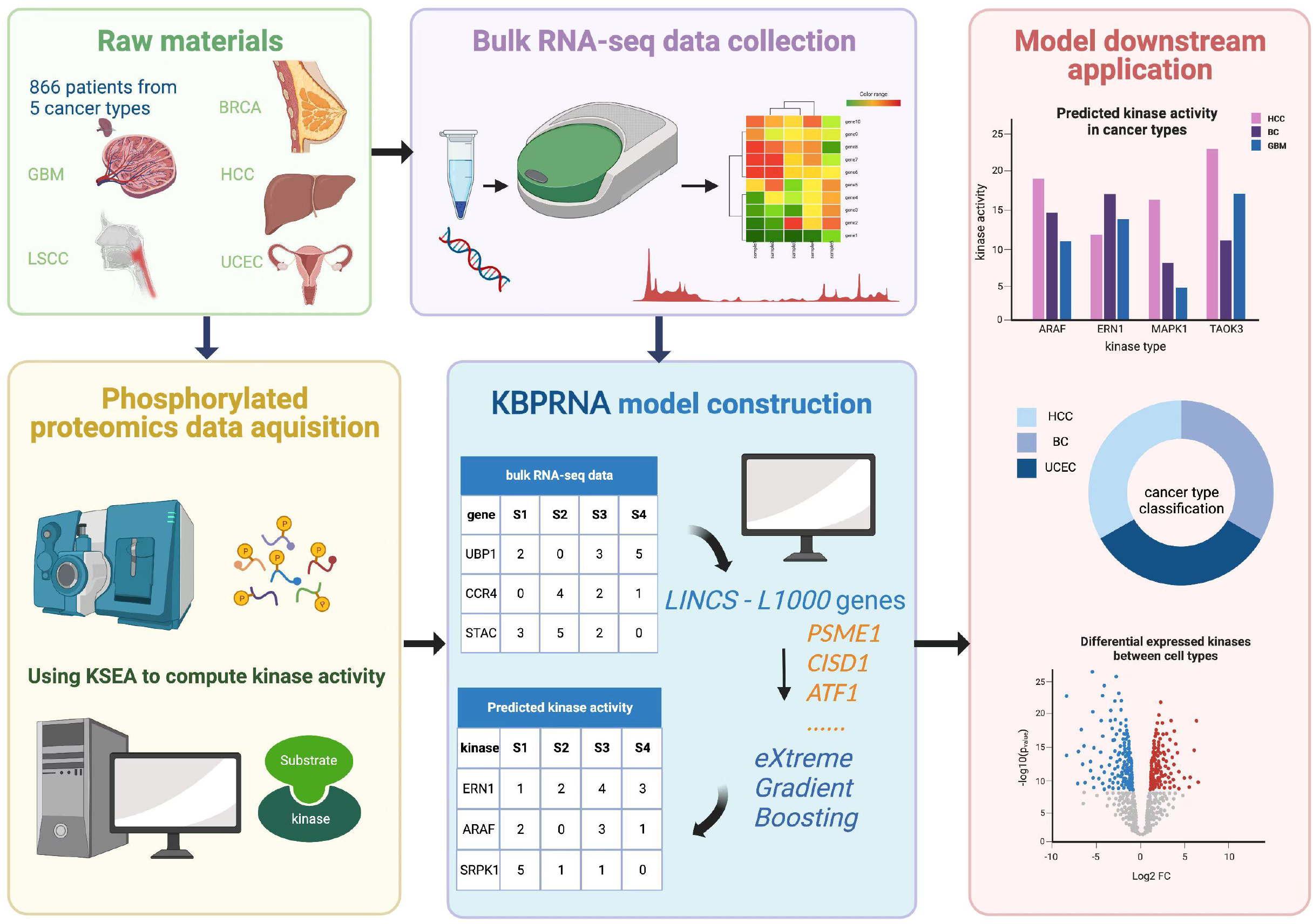

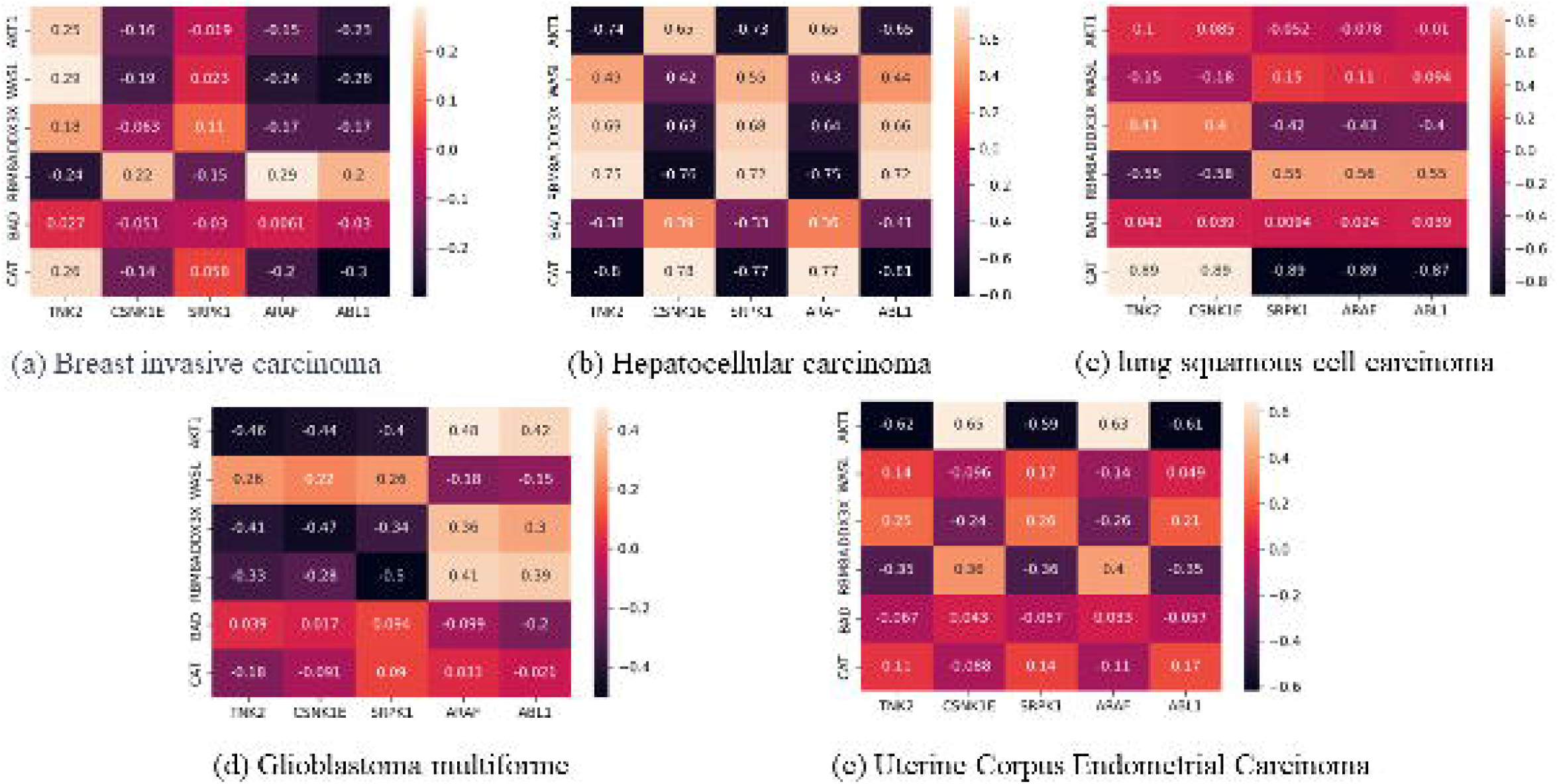

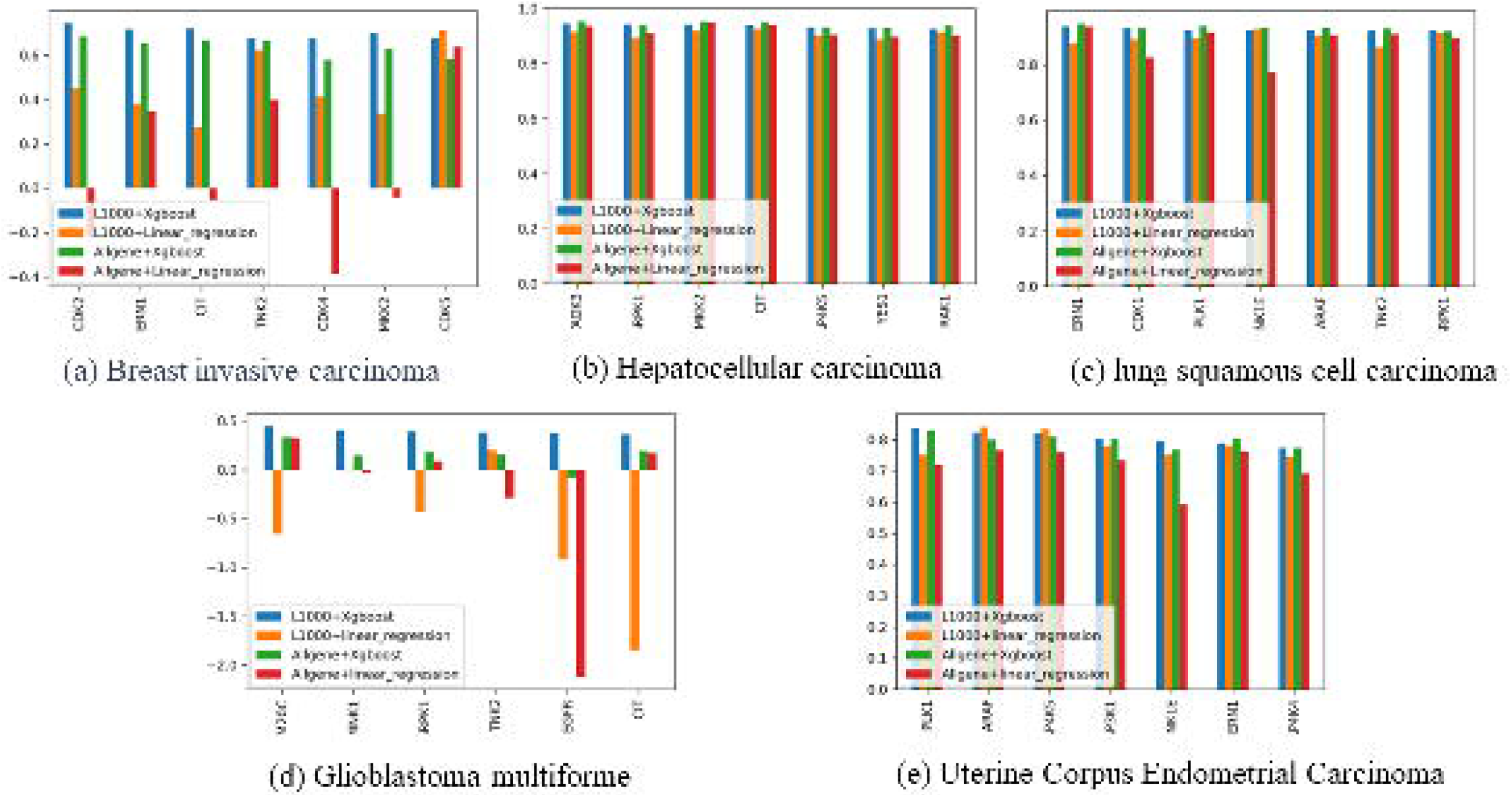

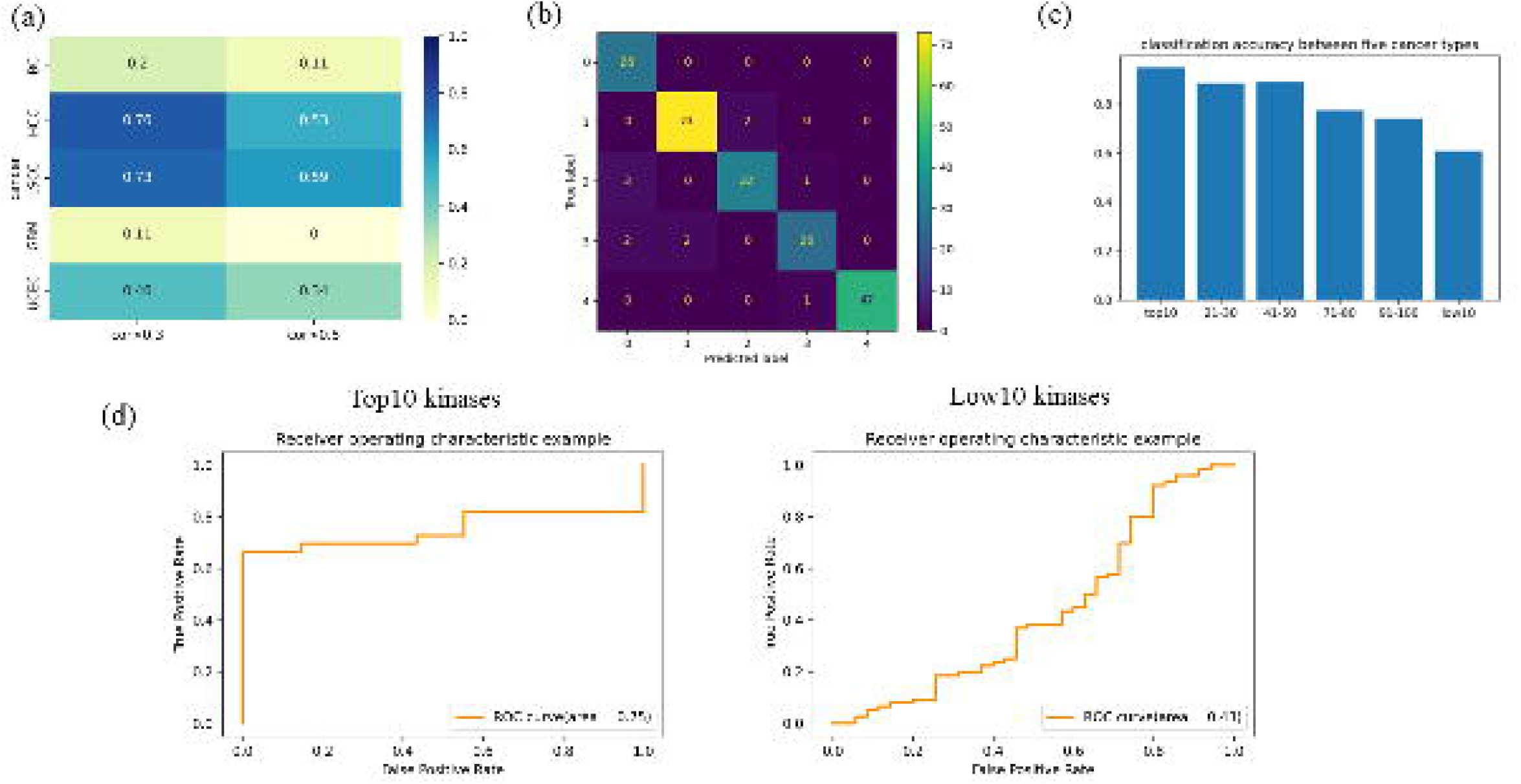

### Application of KBPRNA to scRNA-seq dataset of breast invasive carcinoma tissue

KBPRNA could also be applied to scRNA-seq dataset of cancer tissues to identify novel kinases as biomarkers to differentiate specific cell types. Samples from cancer tissues are usually composed of three cell types which are immune cells, epithelial cells, stromal cells[32]. PTPRC, PECAM1, EPCAM are used as marker genes for immune cells, stromal cells and epithelial cells respectively. After downloading scRNA-seq dataset GSM5457199[25] which comes from tissues of breast invasive carcinoma, we used scanpy to disentangle cell compositions of this sample [33]. We annotated each cell group manually and labeled corresponding cell names on the cell groups (Fig 5B, C). KBPRNA was utilized to construct kinase activity prediction model for 50 kinases with high predictability under condition of breast invasive carcinoma (Fig5D). Heatmap summarizing total 50 kinase activities for four cell groups was shown in Fig5D. This model could be utilized to identify kinases as biomarkers to differentiate various cell groups.

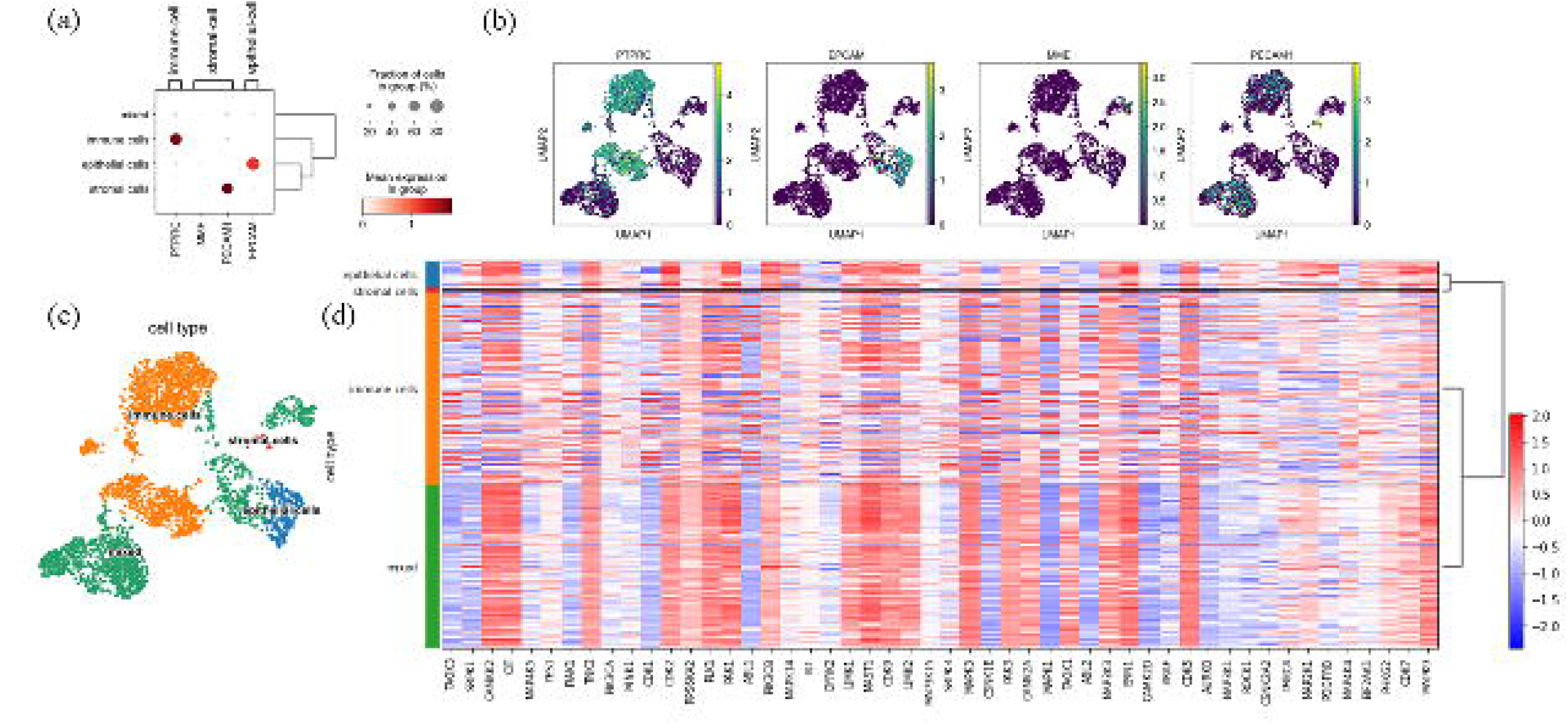

## Discussion

To investigate KBPRNA’s power in predicting kinase activities, we utilized R square as metrics to measure the performance of the model among tissues from five cancer types (breast invasive carcinoma, hepatocellular carcinoma, lung squamous cell carcinoma, glioblastoma multiforme, uterine corpus endometrial carcinoma). According to the prediction accuracy of this model, we validate the model’s robustness and efficiency in obtaining specific and significant kinase activities. Our model achieved better performance than other baseline models in kinase activity prediction from bulk RNA-seq dataset originating from cancer tissues. Then we made a list of kinases through ranking their predictability. Kinases with high predictability were found to be reliable in classifying multiple cancer types. Finally, we produced 50 kinase activity prediction model in order to construct kinase activity profiles from scRNA-seq samples of breast invasive carcinoma. We identified several kinases with significantly differential activities between each cell group. These results prove reliability of KBPRNA and indicate possible downstream usage. Kinase activities have seldom been predicted utilizing bulk RNA-seq dataset coming from human tissues as input features. Here, we introduced a novel machine learning method KBPRNA to construct kinase activity profiles when at least 30%kinases could be predicted accurately with R square over 0.5. In previous research, Sam Crowl et al devised an algorithm called KSTAR to predict patient specific kinase activities from phosphoproteomic data [34]. Our method fills the gap between bulk RNA-seq data and kinase activities instead of achieving phosphoproteomics data in advance from cancer tissues. We built kinase activity prediction model across five cancer types and this model could be utilized to construct kinase activity profiles.

Cancer type identification has been a popular topic in the field of bioinformatics. Lu et al. found miRNA profiles could be used to do cancer classification [35]. Our research validated the usage of kinase activities in cancer type identification. High classification accuracy proved that kinases with high predictability based on the bulk RNA-seq data could be used to classify specific cancer type.

Recently, single cell RNA-seq technology has emerged and really reshaped our knowledge of traditional biology. We would like to identify differentially expressed kinases to differentiate different cell groups. Our model robustly built single cell kinase activity profiles of breast invasive carcinoma and identified several kinases which have the ability to differentiate different cell groups. Our identified kinases could be further experimentally validated in order to be used as clinical markers and promote healthcare improvement.

While our model shows its advantage in cancer type classification and specific kinase activity prediction, there are several drawbacks of our research. Firstly, kinase activity prediction performance may vary from cancer to cancer. Our model performs better in tissues of Hepatocellular carcinoma and lung squamous cell carcinoma than in tissues of Glioblastoma multiforme. Thus, our model needed to be modified to fit for the tissues of Glioblastoma multiforme. Secondly, in order to prove the robustness of our model in advance, more experiments in vivo or in vitro are needed to validate the functions of specific kinases identified by our model. Only in this way can we confirm the functions of kinases we identified. Last but not least, more study with larger datasets will be required in order to determine whether KBPRNA could be successfully used for kinase activity prediction based on bulk RNA-seq datasets from cancer conditions. That measure will improve the robustness of our model and promote more extension of our research.

## Conclusion

For five cancer types we studied,KBPRNA which combines both LINCS-L1000 gene signatures and bulk RNA-seq datasets outperformed other benchmarking methods in kinase activity prediction. This model achieved an accurate estimation of over fifty percent kinases’ activity with R square over 0.5 in tissues of Hepatocellular carcinoma and lung squamous cell carcinoma. Given a raw bulk RNA-seq dataset, this model is a powerful tool in constructing kinase activity profiles of specific cancer types. Apart from been applied to produce kinase activity profiles, KBPRNA could also be used to identify abnormal kinases which contribute to the pathogenesis of specific cancer diseases. Finally, this model could be utilized to rapidly estimate specific kinases’ functions in cancer diseases and aid in clinical prognosis. In the future, we hope to include more cancer tissue samples to improve this model’s applicability and provide a wider understanding of specific kinases’ functions in the pathogenesis of cancer diseases.

## Availability of data and materials

All the code used to prepare the data and fit the models is available at https://github.com/tibettiger/kinase_prediction. The dataset analyses during the current study are published in GEO website.

## References

1. Wang C, Xu H, Lin S, Deng W, Zhou J, et al. GPS 5.0: an update on the prediction of kinase specific phosphorylation sites in proteins. Genom, Proteom & Bioinform. 2020; 18(1): 72–80.

2. Paul MK, Mukhopadhyay AK. Tyrosine kinase–role and significance in cancer. International Journal of Medical Sciences. 2004; 1(2): 101.

3. Zwick E, Bange J, Ullrich A. Receptor tyrosine kinases as targets for anticancer drugs. Trends in Molecular Medicine. 2002; 8(1): 17–23.

4. Roskoski JR. Properties of FDA-approved small molecule protein kinase inhibitors: a 2020 update. Pharmacological Research [Internet]. 2020 [cited 2022 Nov 11]. Available from: https://www.sciencedirect.com/science/article/pii/S1043661819328890

5. Hasinoff BB, Patel D. The lack of target specificity of small molecule anticancer kinase inhibitors is correlated with their ability to damage myocytes in vitro. Toxicology and Applied Pharmacology. 2010; 249(2): 132–139.

6. Casnellie JE, Krebs EG. The use of synthetic peptides for defining the specificity of typrosine protein kinases. Advances in Enzyme Regulation [Internet]. 1984 [cited 2022 Nov 11]. Available from: https://www.sciencedirect.com/science/article/abs/pii/0065257184900281

7. Casnellie JE. Assay of protein kinases using peptides with basic residues for phosphocellulose binding. Methods in Enzymology [Internet]. 1991 [cited 2022 Nov 11]. Available from: https://www.sciencedirect.com/science/article/abs/pii/007668799100133H

8. Wang Y, Ma H. Protein kinase profiling assays: a technology review. Drug Discovery Today: Technologies [Internet]. 2015 [cited 2022 Nov 11]. Available from: https://www.sciencedirect.com/science/article/pii/S1740674915000505

9. Gonzalez-Vera JA. Probing the kinome in real time with fluorescent peptides. Chemical Society Reviews. 2012; 41(5): 1652–1664.

10. Crowl S, Jordan BT, Ahmed H, Ma CX, Naegle KM. KSTAR: an algorithm to predict patient specific kinase activities from phosphoproteomic data. Nature Communications [Internet]. 2022 [cited 2022 Nov 11]. Available from: https://pubmed.ncbi.nlm.nih.gov/35879309/

11. Lo YC, Liu T, Morrissey KM, Kakiuchi-Kiyota S, Johnson AR, et al. Computational analysis of kinase inhibitor selectivity using structural knowledge. Bioinformatics. 2019; 35(2): 235–242.

12. Yang Z, Ye Z, Xiao Y, Hsieh C, Zhang S. SPLDExtraTrees: robust machine learning approach for predicting kinase inhibitor resistance. Briefings in Bioinformatics. 2022; 23(3): 1–12.

13. Fenoy E, Izarzugaza JMG, Jurtz V, Brunak S, Nielsen M. A generic deep convolutional neural network framework for prediction of receptor–ligand interactions—NetPhosPan: application to kinase phosphorylation prediction. Bioinformatics. 2019; 35(7): 1098–1107.

14. Kirchoff KE, Gomez SM. EMBER: multi-label prediction of kinase-substrate phosphorylation events through deep learning. Bioinformatics. 2022; 38(8): 2119–2126.

15. Patrick R, Horin C, Kobe B, Cao KAL, Bodén M. Prediction of kinase-specific phosphorylation sites through an integrative model of protein context and sequence. Biochimica et Biophysica Acta - Proteins Proteom. 2016; 1864(11): 1599–1608.

16. Wang LB, Karpova A, Gritsenko MA, Kyle JE, Cao S, et al. Proteogenomic and metabolomic characterization of human glioblastoma. Cancer Cell. 2021; 39(4): 509–528.

17. Krug K, Jaehnig EJ, Satpathy S, Blumenberg L, Karpova A, et al. Proteogenomic landscape of breast cancer tumorigenesis and targeted therapy. Cell. 2020; 183(5):1436–1456.

18. Charlotte KYN, Dazert E, Boldanova T, Coto-Llerena M, Nuciforo S, et al. Integrative proteogenomic characterization of hepatocellular carcinoma across etiologies and stages. Nature Communications [Internet]. 2022 [cited 2022 Nov 11]. Available from: https://www.nature.com/articles/s41467-022-29960-8

19. Pan L, Wang X, Yang L, Zhao L, Zhai L, et al. Proteomic and phosphoproteomic maps of lung squamous cell carcinoma from Chinese patients. Front Oncol [Internet]. 2020 [cited 2022 Nov 11]. Available from: https://www.frontiersin.org/articles/10.3389/fonc.2020.00963/full

20. Dou Y, Kawaler EA, Zhou DC, Gritsenko MA, Huang C, et al. Proteogenomic characterization of endometrial carcinoma. Cell. 2020; 180(4):729–748.

21. Erhard F, Saliba AE, Lusser A, Toussaint C, Hennig T, et al. Time-resolved single-cell RNA-seq using metabolic RNA labelling. Nature Reviews Methods Primers. 2022; 2(1): 1–18.

22. Perkel JM. The software that powers scientific illustration. Nature. 2020; 582(7810): 137–139.

23. Zhang K, Erkan EP, Jamalzadeh S, Dai J, Andersson N, et al. Longitudinal single-cell RNA-seq analysis reveals stress-promoted chemoresistance in metastatic ovarian cancer. Science Advances [Internet]. 2022 [cited 2022 Nov 11]. Available from: https://www.science.org/doi/full/10.1126/sciadv.abm1831

24. Wiredja DD, Koyutürk M, Chance MR. The KSEA App: a web-based tool for kinase activity inference from quantitative phosphoproteomics. Bioinformatics. 2017; 33(21): 3489–3491.

25. Bai M, Sun C. Determination of breast metabolic phenotypes and their sssociations with immunotherapy and drug-targeted therapy: analysis of single-cell and bulk sequences. Frontiers in Cell and Developmental Biology [Internet]. 2022 [cited 2022 Nov 11]. Available from: https://www.ncbi.nlm.nih.gov/pmc/articles/PMC8905618/

26. Luo Q, Mo S, Xue Y, Zhang X, Gu Y, et al. Novel deep learning-based transcriptome data analysis for drug-drug interaction prediction with an application in diabetes. BMC Bioinformatics. 2021; 22(1): 1–15.

27. Lu J, Chen M, Qin Y. Drug-induced cell viability prediction from LINCS-L1000 through WRFEN-XGBoost algorithm. BMC Bioinformatics. 2021; 22(1): 1–18.

28. Ranjan GSK, Verma AK, Radhika S. K-nearest neighbors and grid search cv based real time fault monitoring system for industries. IEEE 5th International Conference for Convergence in Technology (I2CT) [Internet]. 2019 [cited 2022 Nov 11]. Available from: https://www.researchgate.net/publication/339909406_K-Nearest_Neighbors_and_Grid_Search_CV_Based_Real_Time_Fault_Monitoring_System_for_Industries

29. Jha A, Quesnel-Vallières M, Wang D, Thomas-Tikhonenko A, Lynch KW, et al. Identifying common transcriptome signatures of cancer by interpreting deep learning models. Genome Biology [Internet]. 2022 [cited 2022 Nov 11]. Available from: https://genomebiology.biomedcentral.com/articles/10.1186/s13059-022-02681-3

30. Liu C, Su J, Yang F, Wei K, Ma J, et al. Compound signature detection on LINCS L1000 big data. Molecular BioSystems. 2015; 11(3): 714–722.

31. Santhanam R, Uzir N, Raman S, Banerjee S. Experimenting XGBoost algorithm for prediction and classification of different datasets. International Journal of Control Theory and Applications [Internet]. 2016 [cited 2022 Nov 11]. Available from: https://www.researchgate.net/publication/318132203_Experimenting_XGBoost_Algorithm_for_Prediction_and_Classification_of_Different_Datasets

32. Yoshihara K, Shahmoradgoli M, Martínez E, Vegesna R, Kim H, et al. Inferring tumour purity and stromal and immune cell admixture from expression data. Nature communications. 2013; 4(1): 1–11.

33. Wolf F A, Angerer P, Theis FJ. SCANPY: large-scale single-cell gene expression data analysis. Genome Biology. 2018; 19(1): 1–5.

34. Crowl S, Jordan BT, Ahmed H, Ma CX, Naegle KM. KSTAR: an algorithm to predict patientspecific kinase activities from phosphoproteomic data. Nature Communications. 2022; 13(1): 1–16.

35. Lu J, Getz G, Miska EA, Alvarez-Saavedra E, Lamb J, et al. MicroRNA expression profiles classify human cancers. Nature [Internet]. 2005 [cited 2022 Nov 11]. Available from: https://www.nature.com/articles/nature03702

